# Gelbox — An Interactive Simulation Tool for Gel Electrophoresis

**DOI:** 10.1101/406132

**Authors:** Chaim Gingold, Shawn M. Douglas

## Abstract

Gel electrophoresis enables separation and visualization of biomolecules such as DNA, RNA, or proteins. Like many powerful tools, mastering the use of gels and making the information gained from them productive, can be difficult. Gelbox is a simulation tool that helps users understand how changing experimental input parameters can affect the data output from gel electrophoresis. Our simulation model handles a wide range of settings, including many “suboptimal” values that may be useful for troubleshooting common mistakes made by novices.

## Introduction

In 1948, when Arne Tiselius was awarded Nobel Prize in Chemistry for his pioneering work on electrophoresis, he summarized his motivations for the research as follows^1^:

*[In biochemistry] some of the most important problems are to isolate the substances which are responsible for a specific biological or biochemical effect, and also to define and characterize these substances as accurately as possible.*

Today, in labs worldwide, researchers use gel electrophoresis to isolate, define, and characterize biomolecules. Gel electrophoresis is a powerful and versatile technique that is remarkably precise, yet preserves the structures of the molecules being analyzed. However, for a variety of reasons, the technique can be difficult to master.

First, gels must be prepared according to detailed protocols. Small variations in the amount of an ingredient or the timing of a step can dramatically affect gel behavior. To illustrate this concept, we ran identical 1 kb ladder samples on three agarose gels that were prepared with different buffers:

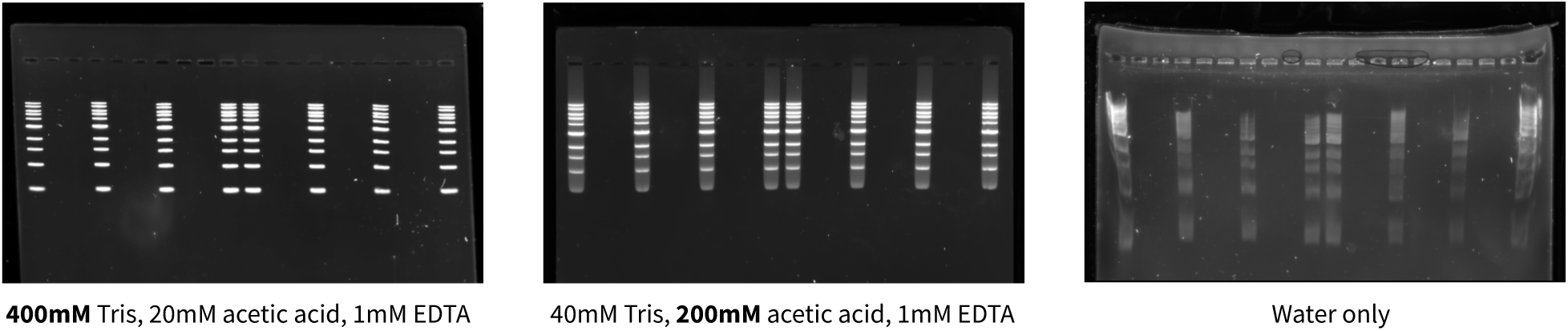

We observed that the degree of band mobility, smearing, and distortion varied greatly for each gel. Even when gels are run successfully, interpreting data can be tricky because the relationships between “band” patterns and the molecules in a sample are abstract and often ambiguous. Challenges in preparing and using gels can frustrate budding scientists, interfere with scientific reproducibility, and impede overall research progress.

We believe interactive simulation and visualization tools can help. We created Gelbox, a dynamic “scientific sandbox” that can aid in visualizing various ways that gel bands, sample molecules, and the gel itself, interact. We wanted to explore the potential pitfalls of using gels by illustrating that, although it’s easy to make mistakes when using this method, these mistakes are also predictable and can mostly be avoided with some training and forethought.

This project builds on concepts from *Earth: A Primer*, an interactive geology science book by Gingold^2^. We have been greatly influenced by Bret Victor’s work, especially *Up and Down the Ladder of Abstraction*^3^. We also drew inspiration from *Parable of the Polygons* by Vi Hart and Nicky Case^4^, and many other works^5–7^.

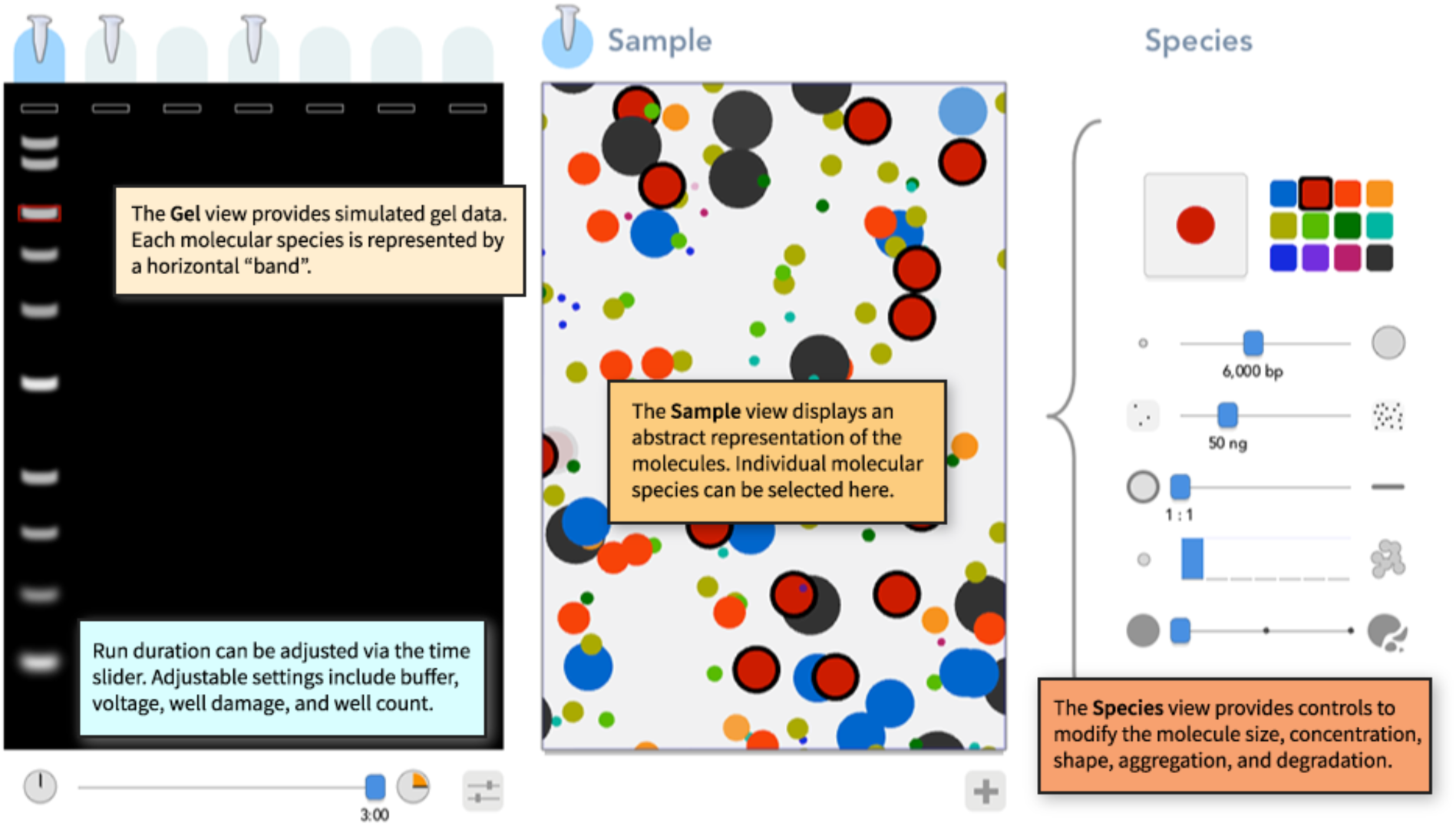

### The Gelbox Interface

### Commands & Shortcuts

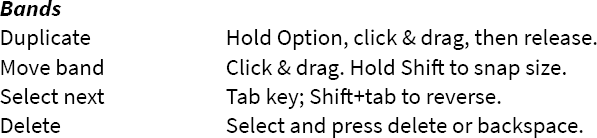
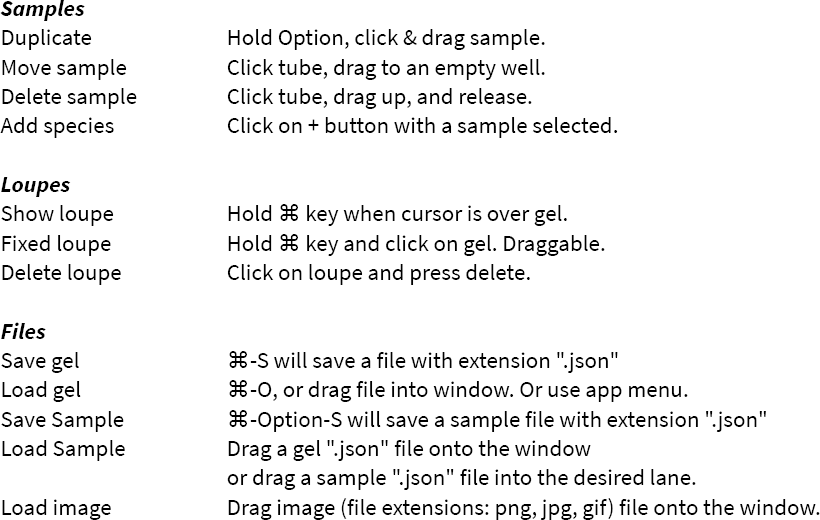

## Discussion

We built Gelbox to be an interactive tool that could complement the *ad hoc* learning methods routinely used in laboratory settings. While working with real gels provides valuable hands-on experience, the learning process is constrained by a slow feedback loop. Several hours can pass between making a mistake in preparing a gel and observing the effects of that mistake. Fortunately, computer simulation can reduce the time to fractions of a second between modifying a gel parameter and observing the effects, making cause-and-effect relationships more visible and explicit.

While other gel simulators do exist, they provide only a single static graphical representation of predicted DNA fragment mobility under ideal experimental conditions^8–11^. These tools are typically included with computer-aided design software for gene editing or DNA cloning. In contrast, Gelbox provides multiple dynamic representations of a gel under a wide range of conditions. Molecules can vary in size, concentration, shape, or degree of aggregation and degradation. To our knowledge, Gelbox is the only gel simulation tool that attempts to capture and explain how gels can produce flawed information, particularly due to user error, or fail completely.

To build the simulation model, we aimed to write code that would accurately convert the length of a DNA fragment to band mobility in the gel. We explored models from Van Winkle, Beheshti, and Rill^12^, and the dissertation of A. Beheshti^13^, but ultimately developed our own model directly from gel-image data for a 1 kb ladder.

Gelbox has several limitations. First, Gelbox only models agarose gels and DNA fragments ranging in size from 100 to 14000 base pairs. It does not yet support RNA or protein gels. Second, the simulation model is narrow in scope compared to real agarose gels. We spent most of our effort on creating the various representations and thus didn’t capture the great variation in possible parameters. Third, the simulation could benefit from further calibration. We did not systematically validate all parameters with experimental data, thus the predicted band mobility and behavior may not match real-world results. Fourth, our models do not have any physical basis: that is, we do not account for kinetic or thermodynamic properties of the DNA molecules. However, Gelbox might be used as a visualization tool for other physically based simulations if they were adapted to output our text-based file format.

In summary, we have created a dynamic, interactive gel electrophoresis simulator that relies on multiple linked representations of the system to help young or new researchers achieve more accurate and useful results when using gel electrophoresis. Gelbox may help learners establish some basic skills to help them interpret gel data and to troubleshoot problematic experimental conditions. Many of the limitations of Gelbox could be addressed with further development, including more experimental validation and integration with other simulation tools. We hope our work inspires others to build interactive simulations for additional aspects of scientific research so that all of these efforts can be synergistic, and so Gelbox becomes more useful and accurate with further iterations.

## Acknowledgements

We thank J Brown for assistance with the app icon and introductory video, and P Nafisi for running several gels that we used to calibrate the simulation model. We thank P Rothemund and N Case for helpful testing and feedback. This work was supported by Pew-Stewart Scholars Program for Cancer Research, the National Science Foundation Center for Cellular Construction (CCF 1317694, DBI 1548297), and the UCSF Program for Breakthrough Biomedical Research, which is partially funded by the Sandler Foundation.

